# Neural Underpinnings of Olfactory Dysfunction across Parkinson’s and Alzheimer’s Spectra

**DOI:** 10.1101/2025.10.07.681050

**Authors:** Zhengui Yang, Hiroki Togo, Mitsunari Abe, Toshiya Murai, Fukiko Kitani-Morii, Kazuaki Kanai, Naoya Oishi, Parkinson’s and Alzheimer’s Disease Dimensional Neuroimaging Initiative (PADNI), Takashi Hanakawa

**Author notes:** Correspondence to: Takashi Hanakawa.

## Abstract

Olfactory dysfunction is a frequent yet understudied feature of neurodegenerative spectrum disorders, including Alzheimer’s disease (AD) and Parkinson’s disease (PD). To disentangle the neural substrates of hyposmia across disease spectra, we examined 222 participants from the Parkinson’s and Alzheimer’s disease Dimensional Neuroimaging Initiative cohort. Participants were classified according to the presence of cognitive disturbance, movement disorder, or both. Olfactory testing disclosed that cognitive disturbance and movement disorder were independently associated with hyposmia, and having both cognitive disturbance and movement disorder was associated with severe hyposmia. Whole-brain voxel-based morphometry revealed that hyposmia was associated with atrophy in the medial temporal lobe (MTL) in individuals with cognitive disturbance, whereas an artificial intelligence-based segmentation model identified olfactory bulb atrophy in those with movement disorder. Regression analysis and structural equation modeling further confirmed that the MTL and olfactory bulb volume contributed to olfactory performance through distinct processes. Individuals with cognitive disturbance and movement disorder had atrophy in both the MTL and olfactory bulb (“double hit”). We identified dual processes underlying hyposmia in the AD and PD spectra: a process linking MTL degeneration to cognitive disturbance and a process linking olfactory bulb degeneration to movement disorder. Our transdiagnostic approach enhances strategy in identifying specific neural correlates underlying hyposmia, lending support to the development of biomarkers for early intervention in AD and PD.

## Introduction

Olfactory dysfunction refers to the impaired perception of smell, which varies from hyposmia, a diminished sensitivity to odours, to anosmia, its complete loss^1^. While olfaction declines with age^2^, it is also under the influence of various pathological processes. Particularly, olfactory dysfunction is recognized as a prodromal or adjunct symptom in both Alzheimer’s Disease (AD) and Parkinson’s Disease (PD), the two most prevalent neurodegenerative disorders^3,4^. Increasing attention has been directed toward utilizing olfaction as a marker for prediction, early detection, and stratification in AD and PD^5–7^.

Odour information is initially processed by the olfactory epithelium, transmitted to the olfactory bulb and to the olfactory cortex, and then to higher-order regions in the medial temporal lobe (MTL)^8–10^. Many studies have examined the relationship between olfactory dysfunction and brain structural alterations. In AD, olfactory dysfunction has consistently been linked to atrophy of the MTL, including the hippocampus, amygdala, and parahippocampal gyrus, which are closely associated with cognition and memory^11–13^. This pattern parallels tau-neurofibrillary tangle (NFT) pathology, which initially accumulates in the MTL and subsequently propagates to the primary olfactory regions^14,15^. Contrarily, the neural correlates of olfactory dysfunction remain unclear in PD, as previous studies reported heterogeneous results. Atrophy of the olfactory bulb^16^ or that of the piriform cortex^17^ was reported, whereas striatal abnormalities are also found^18^. These findings, in part, align with Lewy body propagation patterns, in which the olfactory bulb is among the earliest sites where the pathology starts^19^. These different pathology propagation patterns between AD and PD prompted us to consider two opposing processes: the afferent process, which involves Lewy body pathology initiated in the olfactory bulb, and the efferent process, which involves tau-related degeneration originating from the MTL. Testing this idea requires examining the neural correlates of olfactory dysfunction in sufficiently large samples of AD and PD. Notably, almost all those previous studies have focused exclusively on either AD^11,12^ or PD^17,18,20^. Only one study examined a limited number of patients from both AD and PD, reporting orbitofrontal atrophy correlated with hyposmia in AD, but not in PD^21^. Accordingly, the distinct or shared neural mechanisms underlying olfactory impairment in these two most prevalent neurodegenerative disorders remain elusive.

Here, we examined data from a single cohort, including both AD and PD, and related conditions^22^. Recent studies have proposed that AD and PD may represent continuous disease spectra, in which typical cases can be well characterized by a set of clinical and molecular features, while cases that are difficult to label also exist, e.g., those with co-pathology^23–25^. To circumvent the limitations of the disease label approach^26^ and to capture brain-behaviour relationships more objectively across disease spectra^27^, this study employed a factorial grouping strategy based on hallmark symptoms and molecular markers. Specifically, we first labelled each participant based on the presence of cognitive disturbance, movement disorder, or both, and the label was later validated using molecular markers. We hypothesized that olfactory dysfunction would be associated with MTL atrophy in cognitive disturbance and with olfactory bulb atrophy in movement disorder. We employed whole-brain, voxel-based morphometry (VBM) and atlas-based volumetry for testing the MTL atrophy and developed an original artificial intelligence (AI) model to quantify olfactory bulb volume (OBV). This transdiagnostic, symptom-centred neuroimaging approach uncovered the two epicentres damaging the olfactory system across the AD and PD spectra.

## Materials and Methods

### Participants

We analysed interim datasets from 283 participants registered in the Parkinson’s and Alzheimer’s disease Dimensional Neuroimaging Initiative (PADNI), a multicentre cohort study. PADNI recruited clinically diagnosed patients with the AD spectrum or PD spectrum, along with healthy aged individuals, including potentially prodromal disease states. The AD spectrum included AD and mild cognitive impairment (MCI) due to AD, diagnosed according to the National Institute on Aging–Alzheimer’s Association (NIA-AA) criteria^28^. The PD spectrum included PD with and without cognitive decline, diagnosed according to the MDS criteria^29^, and dementia with Lewy bodies (DLB), diagnosed according to the international DLB Consortium criteria^30^. The PD spectrum also included isolated REM behaviour disorder, recognized as one of the strongest risk factors for PD in the Movement Disorder Society (MDS) research criteria for prodromal PD^31^, according to the International Classification of Sleep Disorders, 3^rd^ edition^32^. Healthy controls (HC) were required to have no history of neurological or psychiatric disorders, intact activities of daily living (ADL), and normal neuropsychological performance. Participants were recruited at four clinical sites: the National Centre of Neurology and Psychiatry, Kyoto University, Kyoto Prefectural University of Medicine, and Fukushima Medical University. The inclusion criteria were as follows: age ≥ 50 years and having a study partner who provided information about the participant’s activities of daily living. In the present study, we did not rely on these original diagnostic categories; instead, participants were subsequently reclassified according to symptom dimensions, as described in the following section. The exclusion criteria were as follows: use of medications affecting dopamine uptake (e.g., selective serotonin reuptake inhibitors and tricyclic antidepressants), neurological or psychiatric disorders other than the AD or PD spectrum (e.g., cerebral infarction and major depressive disorder), allergies to alcohol or iodine, and concurrent plans to participate in clinical trials. This study was approved by the ethics committees of all participating institutes (NCNP A2018-089, KU C1435, KPUM 111x, and FMU 222x). The study procedures adhered to the principles outlined in the Declaration of Helsinki, and written informed consent was obtained from all participants.

### Data acquisition

#### Clinical and neuropsychological assessment

All participants underwent standardized clinical and neuropsychological assessments across the groups. For the present study, we used the MDS-sponsored revision of the Unified Parkinson’s Disease Rating Scale, Part III (MDS-UPDRS-III) score, the Japanese versions of the Montreal Cognitive Assessment (MoCA-J), and the Clinical Dementia Rating global scale (gCDR). They were acquired by a neurologist or an experienced psychologist at each site.

#### Odour Identification Testing

For an olfactory functioning test, we used the Odor Stick Identification Test for Japanese (OSIT-J)^33^. The OSIT-J includes twelve odour items familiar to Japanese people (curry, cooking gas, perfume, Japanese cypress, Indian ink, menthol, sweaty socks, rose, wood, roasted garlic, condensed milk, and Japanese orange) and one odourless control. Participants must choose an answer from six options: four odour names (including the correct answer), “detectable but not recognisable,” and odourless. OSIT-J total score to evaluate olfactory function, with higher scores indicating better olfaction.

#### MRI data acquisition

In the PADNI cohort, T1-weighted and T2-weighted three-dimensional (3D) structural MRI data were acquired on a 3-T Verio Dot/Skyra Fit scanner (Siemens, Erlangen, Germany) at NCNP and Skyra scanners (Siemens, Erlangen, Germany) at KU, KPUM, and FMS, according to the Brain/MINDS harmonization protocol termed HARP^34^. T1-weighted 3D images were obtained using the magnetization-prepared rapid gradient-echo (MPRAGE) sequence with the following parameters: repetition time = 2500 ms, echo time = 2.18 ms, inversion time = 1000 ms, flip angle = 8°, field of view = 256 × 240 mm³, matrix size = 320 × 300, and voxel size = 0.8 × 0.8 × 0.8 mm³. T2-weighted 3D images were acquired using a Sampling Perfection with Application optimized Contrasts using different flip angle Evolution (SPACE) sequence with the following parameters: repetition time = 3200 ms, echo time = 565 ms, flip angle = 120°, field of view = 256 × 240 mm², matrix size = 320 × 300, and voxel size = 0.8 × 0.8 × 0.8 mm³.

#### Definition of groups

The original dataset consisted of 283 participants. After excluding participants with missing scale data (either OSIT-J, UPDRS, or gCDR) and those with poor imaging quality, data from 243 participants were available for further assessment. In addition, participants were evaluated using both dopamine transporter (DAT) and amyloid-β (Aβ) imaging results. Twenty-one participants, without cognitive disturbance or movement disorder, were excluded due to abnormal DAT or Aβ PET findings, leaving 222 participants included in this study. MDS-UPDRS part III was used to define participants with significant movement disorder (M+: UPDRS III ≥9) or those without (M−: UPDRS III < 9)^35^, and gCDR was used to define participants with significant cognitive disturbance (C+: gCDR ≥0.5) or those without (C−: gCDR = 0). We thereby divided all the participants into four groups: C+M+ (n = 36), C−M+ (n = 57), C+M− (n = 61), and C−M− (n = 68).

### Whole-Brain Morphometric Analysis

Gray matter volume (GMV) was analysed using a VBM algorithm provided by the Computational Anatomy Toolbox (CAT12, Gaser C, Jena University Hospital, http://dbm.neuro.uni-jena.de/cat/) and SPM12 (Statistical Parametric Mapping, Wellcome Department of Imaging Neuroscience, London, UK). All T1-weighted images were spatially normalized to the Montreal Neurological Institute (MNI) template space, created based on our data, using the Geodesic Shooting (GS) algorithm^36^, and the brain was segmented into grey matter (GM), white matter (WM), and cerebrospinal fluid (CSF). Non-linear warping was then applied to the individual’s GM image to transform it to the GM template in the MNI space. The spatially normalized GM maps were modulated by the Jacobian determinant of the deformation field and corrected for individual brain sizes. The modulated, normalized GM images (voxel size 1.5 × 1.5 × 1.5 mm^3^) were smoothed using an 8-mm full-width at half-maximum isotropic Gaussian kernel. The total intracranial volume (TIV) was computed by summing the GM, WM, and CSF volumes. To address multicentre effects, the ComBat harmonization method was performed at each GM voxel (see details in Supplementary Methods)^37^.

### Olfactory Bulb Volume

Recent advances in high-resolution MRI have enabled the in vivo quantification of olfactory bulb volume^38^. However, since the olfactory bulbs are not covered in the standard whole-brain templates, traditional VBM approaches are not applicable for analysing this structure. To address this limitation, we initially used a previously published method for automated olfactory bulb segmentation on T2-weighted MRI^38^. However, the segmentation quality was suboptimal (see details in Supplementary Methods), prompting us to develop a customized segmentation model tailored to our T2-weighted MRI data. Olfactory bulb segmentation was performed in all 283 participants, as this procedure relies solely on T2-weighted structural images. The model employed a residual 3D U-Net (Supplementary Figure 3) architecture to enhance feature representation and improve training stability. 3D U-Net was trained using manually annotated olfactory bulb images. Each original T2-weighted MRI was cropped to an input image with a size of 64 × 64 × 64 voxels centred at the olfactory bulbs. The model produced probability maps for the left and right olfactory bulbs. Final segmentation masks were obtained by thresholding and connected component filtering. The olfactory bulb volumes (OBVs) were calculated by summing the voxels within the bilateral segmentation masks and were converted to cubic millimetres (mm³). All AI-generated masks were converted back to the native space and visually inspected to ensure quality. The final OBV values used for analysis were calculated as the average across the five cross-validation folds to enhance generalizability and robustness (see the Supplementary Methods for details). To account for the potential site effects, OBV values were harmonized across the sites, using ComBat (see details in Supplementary Methods).

### Statistical analysis

All statistical analyses for demographic data were performed with R version 4.4.1 (R Core Team, 2024). As the primary analysis for olfactory performance, a 2 × 2 factorial analysis of covariance (ANCOVA) was performed to explore the main and interaction effects of motor function (M+ or M−) and cognition (C+ or C−) on olfactory performance, with age included as a covariate. A one-way analysis of variance (ANOVA) and chi-square tests were conducted to explore differences among C+M+, C−M+, C+M−, and C−M− groups in demographic and clinical variables. When necessary, a confirmatory one-way ANCOVA was conducted to control for age and other confounding factors, and *post hoc* comparisons were adjusted for multiple comparisons using the Tukey Honestly Significant Difference (HSD) test. Statistical significance for all tests was set at *P* < 0.05.

VBM multiple regression analyses were conducted to investigate the correlation between regional GMV and OSIT-J scores, controlling for age, gender, TIV, and category labels as nuisance covariates. We employed a statistical threshold of *P <* 0.001 combined with a cluster-level threshold of *P* < 0.05 family-wise error (FWE)-corrected for multiple comparisons. To derive a composite measure of medial temporal lobe GMV (MTL-GMV), GMV was extracted from the olfactory cortex, hippocampus, parahippocampal gyrus, and amygdala, using the Automated Anatomical Labeling (AAL) atlas in standard space and expressed in cubic centimetres (cm^3^). Partial correlation analyses were conducted to examine the associations of MTL-GMV and OBV with behavioural measures, including OSIT-J, MoCA-J, and UPDRS-III. These analyses controlled for age, gender, and TIV. To compare MTL-GMV and OBV across the groups, ANCOVA was conducted, with age, gender, and TIV included as covariates to account for potential confounding effects.

To integrate the findings from VBM and OBV analysis, linear regression models were constructed to assess the relative contributions of regional GMV and OBV to OSIT-J scores, controlling for age, gender, TIV, and disease labels. These values were then summed across hemispheres, yielding a single MTL-GMV value. All continuous predictors were z-standardized prior to analysis. To account for individual differences in brain size and reduce model complexity, both temporal lobe volume and OBV were normalized by TIV. Multicollinearity among factors was assessed using variance inflation factors (VIF). To identify the most parsimonious set of predictors, a backward stepwise regression procedure was applied using the Akaike Information Criterion (AIC) as the criterion for model selection.

To further explore the potential mechanistic pathways, structural equation modelling (SEM) was constructed using the ‘lavaan’ package in R (version 0.6-19)^39^. Model estimation was performed using maximum likelihood with robust standard errors (MLR). To reduce scale-related variance and improve model estimation, all observed variables were standardized using z-scoring in R. Model fit was evaluated using standard indices, including χ², Root Mean Square Error of Approximation (RMSEA), Comparative Fit Index (CFI), Tucker–Lewis Index (TLI), and Standardized Root Mean Square Residual (SRMR). Acceptable model fit was defined as RMSEA < 0.08, CFI > 0.90, and SRMR < 0.08.

## Results

### Demographic and clinical variables

Age differed across the groups (Table 1), depending on the cognitive disturbance factor (C main effect, *F* (1, 218) = 47.75, *P* < 0.001), but not on the M factor (*F* (1, 218) = 0.39, *P* = 0.84) or C × M interactions (*F* (1, 218) = 0.66, *P* = 0.42). This was because the two C+ groups were significantly older than the normal cognition groups (*F* (3, 218) = 17.60, *P* < 0.001 by one-way ANOVA followed by post hoc tests). Gender distribution and TIV did not differ across the groups. A factorial ANCOVA on MoCA-J scores, using age as a covariate, supported the gCDR-defined C+ grouping (Supplementary Table 1). However, MoCA-J scores showed a C × M interaction (*F* (1, 217) = 11.32, *P* < 0.001), mainly because the C+M− group had lower MoCA-J scores than the others (see Table 1). UPDRS-III scores after controlling for age supported the M+ grouping. Molecular profiling also supported the groups (Supplementary Figure 6). Although a detailed correspondence between the clinical and molecular classifications is beyond the scope of this study, the two cognitively disturbed groups exhibited Aβ deposition in the brain, while the two movement disorder groups showed a loss of nigrostriatal dopamine terminals. Together, these findings indicated that the C+M+ group corresponded to DLB or PD-MCI/PD dementia that may accompany Aβ deposition^40^, the C−M+ group to cognitively normal PD, the C+M− group to AD or MCI due to AD, and the C−M− group to normative aging.

**Table 1.**
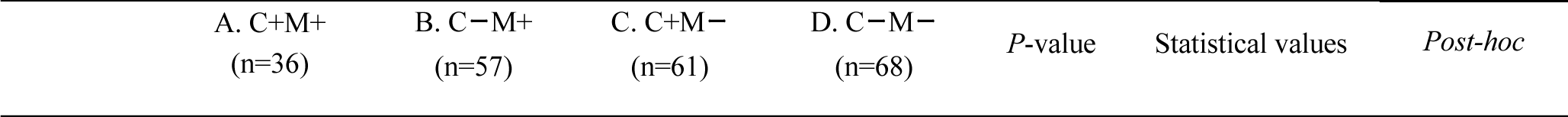
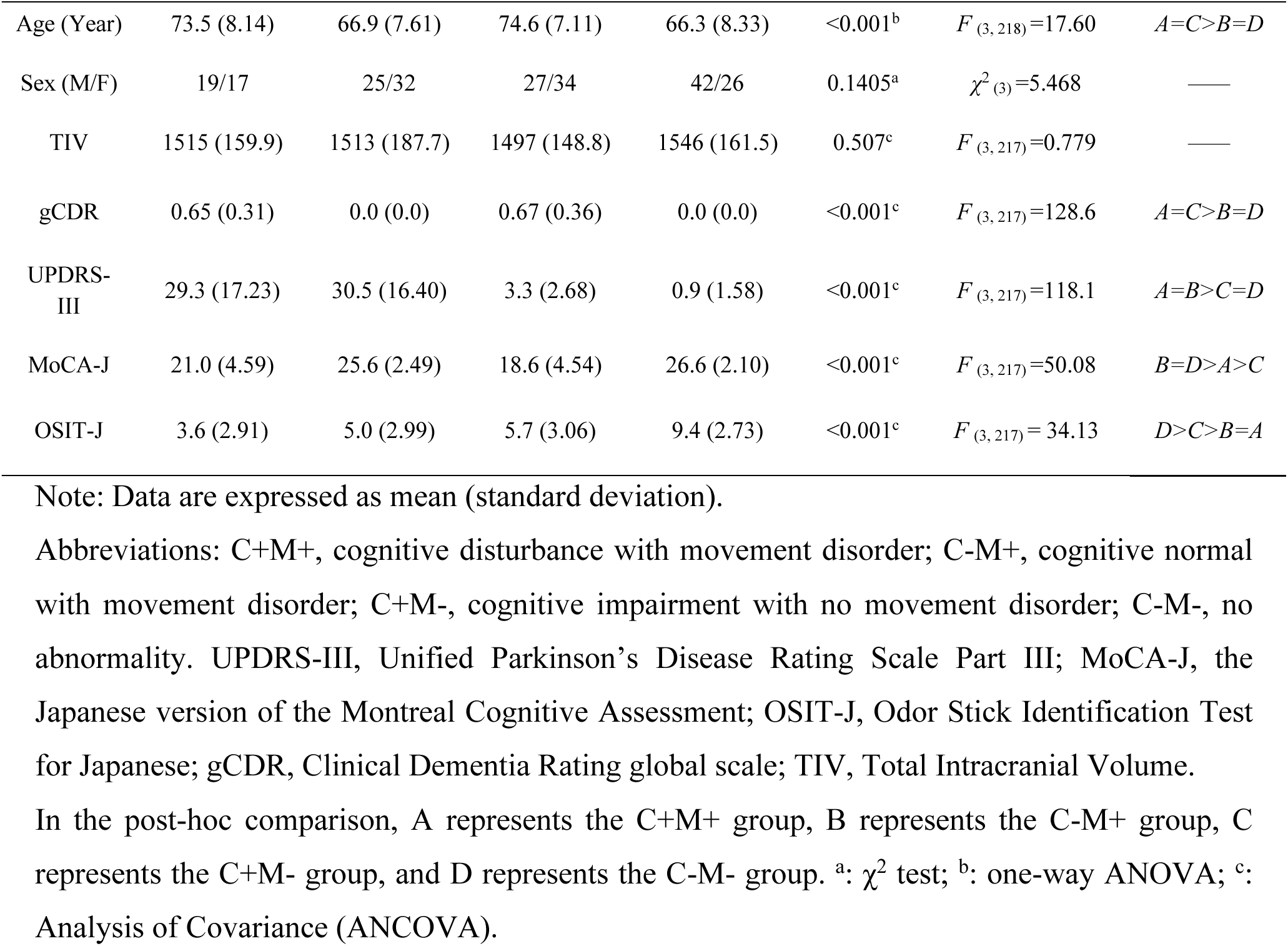
Demographic and neuropsychological characteristics.

| A. C+M+<br>(n=36) | B. C−M+<br>(n=57) | C. C+M−<br>(n=61) | D. C−M−<br>(n=68) | P-value | Statistical values | Post-hoc |
| --- | --- | --- | --- | --- | --- | --- |

|  |  |  |  |  |  |  |  |
| --- | --- | --- | --- | --- | --- | --- | --- |
| Age (Year) | 73.5 (8.14) | 66.9 (7.61) | 74.6 (7.11) | 66.3 (8.33) | <0.001<sup>b</sup> | $F_{(3, 218)}=17.60$ | $A=C>B=D$ |
| Sex (M/F) | 19/17 | 25/32 | 27/34 | 42/26 | 0.1405<sup>a</sup> | $\chi^2_{(3)}=5.468$ | — |
| TIV | 1515 (159.9) | 1513 (187.7) | 1497 (148.8) | 1546 (161.5) | 0.507<sup>c</sup> | $F_{(3, 217)}=0.779$ | — |
| gCDR | 0.65 (0.31) | 0.0 (0.0) | 0.67 (0.36) | 0.0 (0.0) | <0.001<sup>c</sup> | $F_{(3, 217)}=128.6$ | $A=C>B=D$ |
| UPDRS-III | 29.3 (17.23) | 30.5 (16.40) | 3.3 (2.68) | 0.9 (1.58) | <0.001<sup>c</sup> | $F_{(3, 217)}=118.1$ | $A=B>C=D$ |
| MoCA-J | 21.0 (4.59) | 25.6 (2.49) | 18.6 (4.54) | 26.6 (2.10) | <0.001<sup>c</sup> | $F_{(3, 217)}=50.08$ | $B=D>A>C$ |
| OSIT-J | 3.6 (2.91) | 5.0 (2.99) | 5.7 (3.06) | 9.4 (2.73) | <0.001<sup>c</sup> | $F_{(3, 217)}=34.13$ | $D>C>B=A$ |

Note: Data are expressed as mean (standard deviation).
Abbreviations: C+M+, cognitive disturbance with movement disorder; C-M+, cognitive normal with movement disorder; C+M-, cognitive impairment with no movement disorder; C-M-, no abnormality. UPDRS-III, Unified Parkinson's Disease Rating Scale Part III; MoCA-J, the Japanese version of the Montreal Cognitive Assessment; OSIT-J, Odor Stick Identification Test for Japanese; gCDR, Clinical Dementia Rating global scale; TIV, Total Intracranial Volume.
In the post-hoc comparison, A represents the C+M+ group, B represents the C-M+ group, C represents the C+M- group, and D represents the C-M- group. <sup>a</sup>: $\chi^2$ test; <sup>b</sup>: one-way ANOVA; <sup>c</sup>: Analysis of Covariance (ANCOVA).

### Olfactory dysfunctions

In the analysis of OSIT-J scores, participants with cognitive disturbance exhibited more severe hyposmia than those without (C main effects, *F* (1, 217) = 16.86, *P* < 0.001 by factorial ANCOVA). This was also the case for those having movement disorder (M main effects, *F* (1, 217) = 70.58, *P* < 0.001) (Figure 1). The C × M interaction was also significant (*F* (1, 217) = 7.37, *P* = 0.007). The C+M+ group exhibited the most severe hyposmia. One-way ANCOVA followed by post hoc tests revealed that compared to the C−M− group, the other three groups had significantly lower OSIT-J scores. Moreover, the C+M+ (*t* = −3.73, *P* = 0.0013, *d* = −0.78) and C−M+ (*t* = −2.73, *P* = 0.035, *d* = −0.53) groups had more severe hyposmia than the C+M− group (Supplementary Table 2). These results have two major implications: (1) hyposmia coexisted with cognitive disturbance and even more so with movement disorder, and (2) mechanisms underlying hyposmia were distinct between movement disorder and cognitive disturbance^41^, and they might be additive.

**Figure 1.**
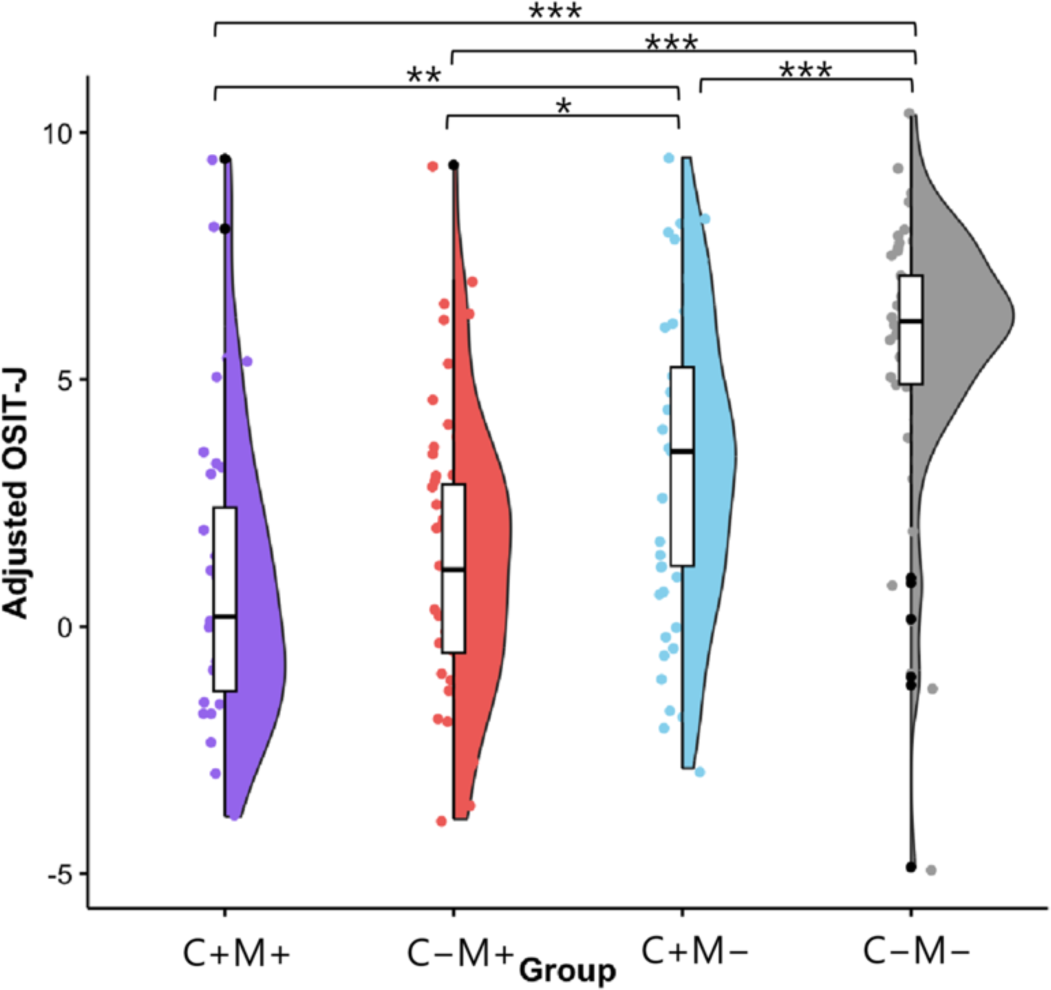
OSIT-J score distribution across groups. The raincloud plot of the Odor Stick Identification Test for Japanese (OSIT-J) scores illustrates the distribution and variation in olfactory functioning among the groups. Significance levels are denoted as follows: *P* < 0.05 (*), *P* < 0.01 (**), *P* < 0.001 (***). Abbreviations: C+M+, cognitive disturbance with movement disorder; C–M+, cognitively normal with movement disorder; C+M–, cognitive disturbance without movement disorder; C–M–, cognitively normal without movement disorder.

### Correlation between GMV and olfaction in all participants

We performed VBM analysis across all participants, with the group label, age, gender, and TIV as covariates. The results showed that hyposmia, as indexed by OSIT-J scores, was correlated with atrophy in the MTL (the amygdala, olfactory cortex, parahippocampal gyrus, and hippocampus) and the middle cingulate cortex (MCC), bilaterally (Figure 2A, Table 2). No brain areas revealed compensatory increases in GMV negatively correlated with OSIT-J scores.

We extracted GMV values from the bilateral amygdala, olfactory cortex, parahippocampal gyri, and hippocampi according to the AAL atlas, and combined them into a single MTL-GMV value. Similarly, we computed MCC-GMV values (Figure 2B). A partial correlation analysis controlling for age, gender, and TIV revealed that both MTL-GMV (*r* = 0.282, *P* < 0.001) and MCC-GMV (*r* = 0.219, *P* = 0.001) were positively correlated with OSIT-J scores, indicating that smaller GMVs in these regions were associated with poorer olfactory performance (Figure 2C). To examine group-wise differences in atrophy patterns, we conducted a factorial ANCOVA on these values, controlling for age, gender, and TIV. Participants with cognitive disturbances showed significantly atrophic MTL compared to those without (main effect of C, *F*(1, 215) = 69.43, *P* < 0.001), whereas the main effect of M was insignificant (*F*(1, 215) = 0.0002, *P* = 0.99). Notably, C × M interaction emerged from this analysis (*F* (1,215) = 10.16, *P* = 0.002). One-way ANCOVA (*F*(3, 215) = 29.62, *P* < 0.001) and *post hoc* tests revealed that the MTL-GMV was especially smaller in the C+ groups compared to the normal cognition groups (*P* < 0.01; Figure 2D and Supplementary Table 3 for details). The C × M interaction was partly accounted for by a tendency toward more severe MTL atrophy in the C+M− group than the C+M+ group (*P* = 0.20, *d* = 0.42, 95% CI [0.005, 0.83]). The smallest MTL of the C+M− group agrees that it had the lowest MoCA-J score. Factorial ANCOVA on MCC-GMV similarly revealed C main effects (*F* (1, 215) = 51.54, *P* < 0.001) and C × M interactions (*F* (1, 215) = 5.20, *P* = 0.02), but no M main effects (*F* (1, 215) = 0.25, *P* = 0.62). One-way ANCOVA (*F* (3, 215) = 20.89, *P* < 0.001) and *post hoc* tests revealed a reduced MCC-GMV in the C+ groups compared with the C− groups (*P* < 0.01; Figure 2D and Supplementary Table 4 for details). Overall, the VBM and atlas-based GMV analyses revealed the involvement of the MTL and MCC in hyposmia among the cognitive disturbance groups, but not in the pure movement disorder group, despite more pronounced hyposmia associated with movement disorder.

**Table 2.**
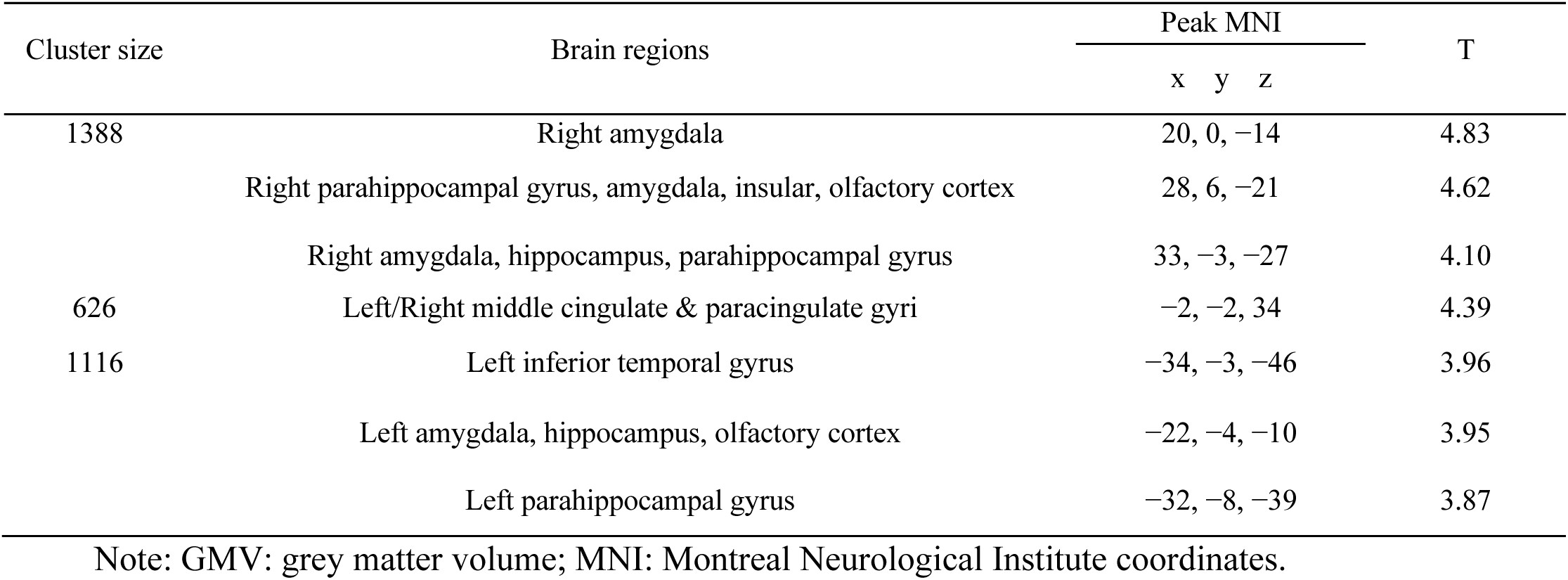
Correlation of olfaction with regional brain GMV.

**Figure 2.**
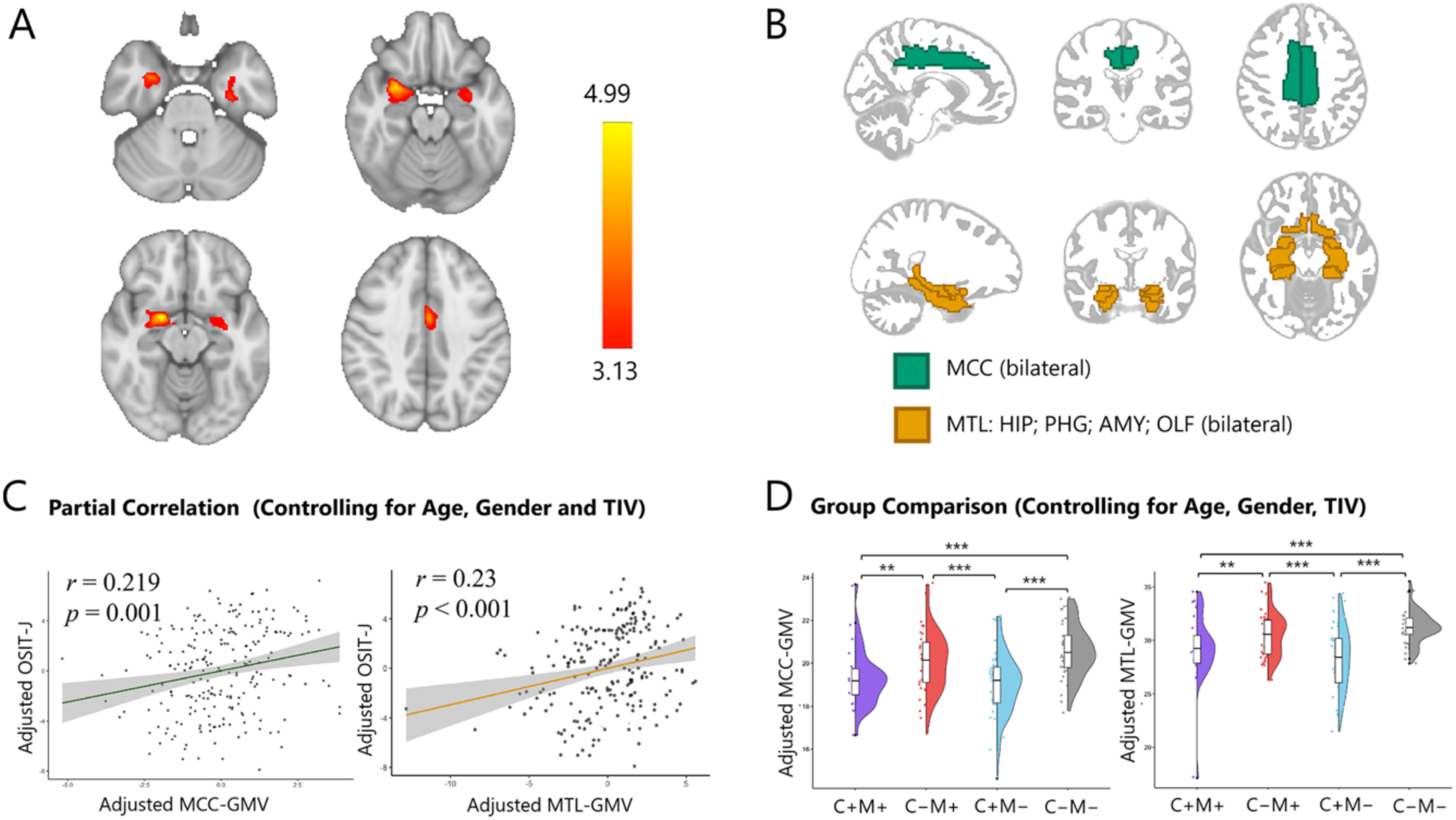
Associations between olfactory function and regional gray matter volume. (**A**) Brain regions associated with olfaction in all participants. (**B**) MCC (amber) was defined on AAL and corresponds to the bilateral middle cingulate cortex; MTL (teal) was defined on AAL as the union of the bilateral Hippocampus (HIP), Parahippocampal gyrus (PHG), Amygdala (AMY) and Olfactory cortex (OLF). (**C**) The partial correlation analysis showing a significant positive association of both MCC-GMV and MTL-GMV with olfactory function (OSIT-J scores). (D) The raincloud plots showing the group-wise comparisons of GMV in the MCC and MTL, with values extracted from the corresponding regions defined by the AAL atlas template. The results remained unchanged when an outlier in the M+C+ group was excluded. Significance levels are denoted as follows: *P* < 0.01 (**), *P <* 0.001 (***). Abbreviations: C+M+, cognitive impairment with movement disorder; C–M+, cognitive normal with movement disorder; C+M–, cognitive impairment with motor normal; C–M–, cognitive normal with motor normal; GMV, grey matter volume; MCC, middle cingulate cortex; MTL, middle temporal lobe.

### Olfactory bulb segmentation and volume measurement

We measured OBV using the residual 3D U-Net model, which generated olfactory bulb masks comparable to those of manually annotated (Figure 3A), with an average Dice score of 0.80 ± 0.038 (as determined by five-fold cross-validation). The out-of-sample validation yielded a Dice score of 0.83 ± 0.02 (Supplementary Figure 7), supporting the model’s generalizability. A group-level consistency map of the olfactory bulb (Figure 3B) confirmed the reliability of the segmentation across participants. The generated masks were used to compute the left and right OBV, which were summed to produce the total OBV used for further analysis.

A partial correlation analysis controlling for age, gender, and TIV revealed a significant positive association between OBV and OSIT-J scores across all participants (*r* = 0.213, *P* = 0.001; Figure 4A). To examine group-specific patterns of OBV changes, a factorial ANCOVA, including age, gender, and TIV as covariates, demonstrated significantly atrophied olfactory bulbs in participants with movement disorder than those without (M main effects, *F* (1, 215) = 13.24, *P* < 0.001). There was no main effect of cognitive disturbance (*F* (1, 215) = 0.08, *P* = 0.78) or a C × M interaction (*F* (1, 215) = 3.21, *P* = 0.07). A one-way ANCOVA followed by a *post hoc* test showed that the M-only (C−M+) group had a smaller total OBV than the C-only (C+M−; *t* = −2.78, *P* = 0.030, *d* = −0.55) and the healthy (C−M−; *t* = −4.155, *P* < 0.001, *d* = −0.75 ) groups (Figure 4B). Although no statistical differences were observed between the other pairs, the C+M+ group tended to have smaller OBV than the C+M− group (*d* = −0.25, 95% CI [−0.66, 0.16]) and the healthy group (*d* = −0.46, 95% CI [−0.88, −0.04]), with small-to-medium effect sizes. To summarize, olfactory bulb atrophy explained hyposmia in the movement disorder groups, but not in the cognitive disturbance groups.

**Figure 3.**
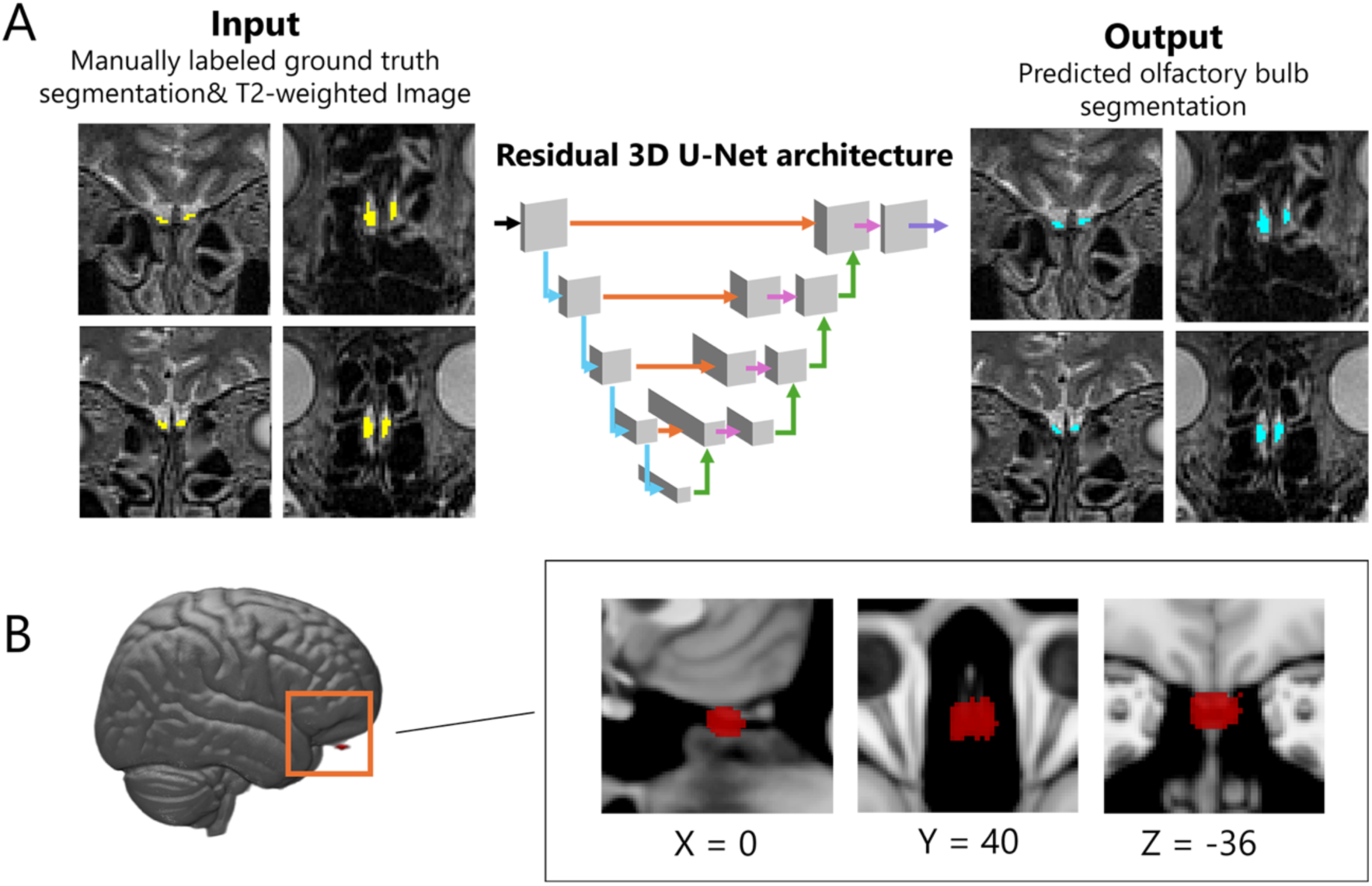
Deep learning–based olfactory bulb segmentation and group-level consistency map. (**A**) Residual 3D U-Net architecture for olfactory bulb segmentation. The network follows an encoder–decoder structure with residual and skip connections. Black arrows indicate residual blocks; light blue arrows represent residual blocks in the encoder; pink arrows represent residual blocks in the decoder; orange arrows denote skip connections; green arrows indicate 3D up-sampling operations; and purple arrows represent 3D convolutional blocks. Manually labeled ground truth segmentation (yellow); Predicted olfactory bulb segmentation (blue); (**B**) Group-level consistency map of olfactory bulb based on T2-weighted segmentation from all participants (n = 283, threshold = 0.02). Red regions indicate voxels identified as part of the olfactory bulb in at least 2% of the participants. Results are overlaid on the standard MNI152 T1-weighted template. The displayed slices correspond to the peak centroid coordinate of the probability map (MNI: x = 0, y = 40, z = –36).

**Figure 4.**
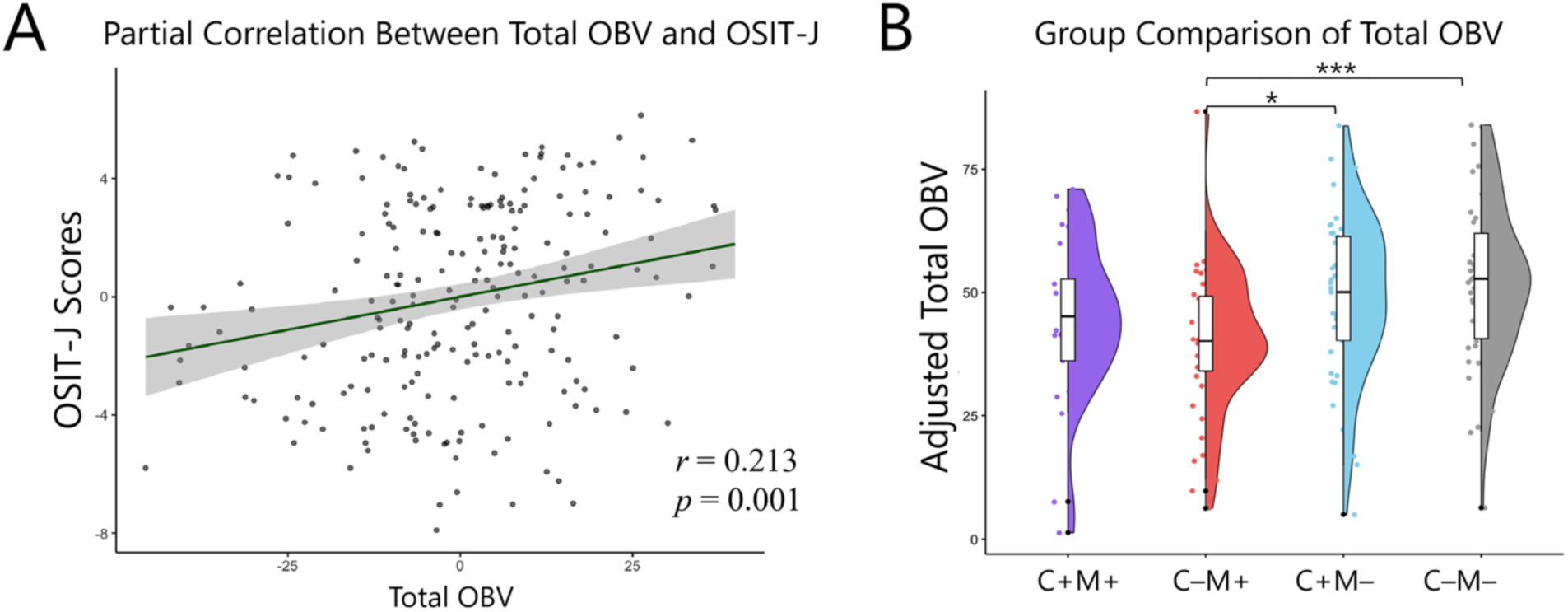
OBV analysis across clinical groups and its clinical correlations. (**A**) Partial correlation analysis showed a significant positive association between OBV and olfactory function (OSIT-J scores); (**B**) Group comparisons of OBV using one-way ANCOVA revealed that the motor-impaired only group (C-M+) showed significantly smaller OBV than both the cognitively impaired only group (C+M−) and healthy elderly controls (C−M−). Significance levels are denoted as follows: *P* < 0.05 (*), *P* < 0.001 (***). Abbreviations: C+M+, cognitive impairment with movement disorder; C–M+, cognitive normal with movement disorder; C+M–, cognitive impairment with motor normal; C–M–, cognitive normal with motor normal. All analyses were adjusted for age, gender, and TIV.

### Liner Model Predicting Olfactory Function

To integrate the findings from the brain GMV and OBV analyses, we conducted a stepwise linear regression analysis that included all participants. There was no substantial multicollinearity among the predictors (VIF < 3). The best model retained age, group labels, MTL-GMV, and total OBV as predictors of OSIT-J scores (AIC = –112.07, *F* (6, 215) = 27.11, *P* < 0.001). This model explained approximately 40% of the variance in olfactory functioning across the participants (adjusted *R^2^* = 0.415).

### Structural Equation Model of Structure-Symptom Relationship

To gain a more holistic understanding of the relationship among demographic and behavioural factors and the underlying neural architecture, we constructed an SEM analysis. Before constructing the model, we examined correlations among clinical and imaging parameters. The OSIT-J scores were positively correlated with MoCA-J scores, MTL-GMV, and total OBV, and negatively correlated with UPDRS-III scores (Supplementary Figure 8).

We thus built the SEM model based on the correlation between the parameters and anatomical connections (Figure 5). The model demonstrated a good fit (*χ²* = 7.938, df = 6, *P* = 0.243; CFI = 0.994; TLI = 0.981; RMSEA = 0.038; SRMR = 0.032). Both MTL-GMV (*β* = 0.296, *P* < 0.001) and total OBV (*β* = 0.219, *P* < 0.001) exerted direct effects on OSIT-J scores. Age negatively predicted OBV (*β* = −0.154, *P* = 0.021) and MTL-GMV (*β* = −0.558, *P* < 0.001). In addition, MTL-GMV predicted MoCA-J (*β* = 0.628, *P* < 0.001). Residual correlations were significant between OSIT-J and UPDRS-III (*r* = −0.390, *P* < 0.001), between OBV and UPDRS-III (*r* = −0.211, *P* = 0.002), and were marginal between OSIT-J and MoCA (*r* = 0.082, *P* = 0.059). These results suggest additional shared variance beyond the specified paths, including the shared neurodegenerative mechanisms that affect both the olfactory and motor systems. Notably, an alternative model including a link between OBV and MTL-GMV did not improve the fit (Supplementary Table 5).

**Figure 5.**
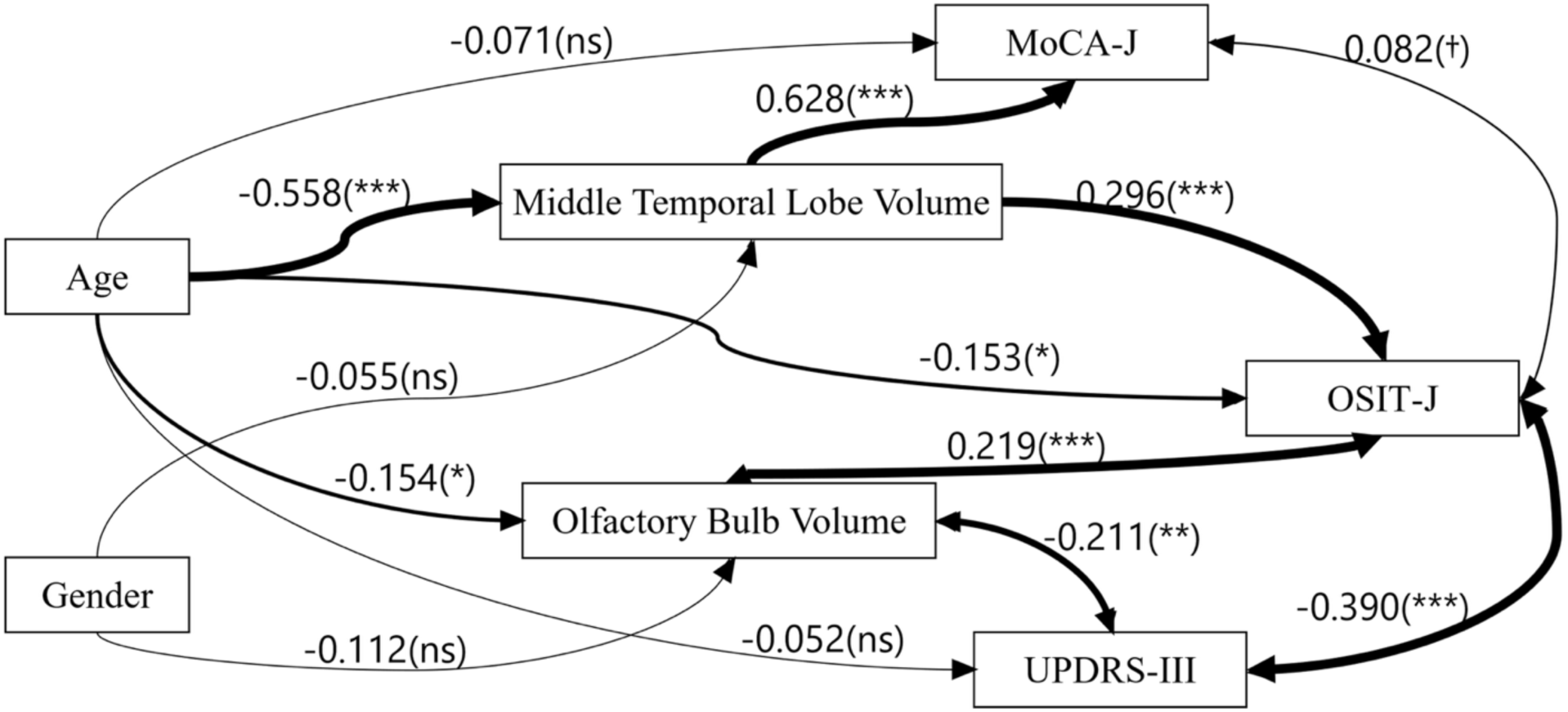
Structural Equation Modeling (SEM) results. Single-headed arrows represent standardized regression coefficients (β), and double-headed arrows represent residual correlations (r). Significance levels are denoted as follows: *P <* 0.05 (*), *P <* 0.01 (**), *P <* 0.001 (***); *P <* 0.10 (†) = marginally significant; “ns” = not significant.

## Discussion

To address the correlates of hyposmia across the AD and PD spectra, we utilized the interim PADNI dataset, which comprises patients with the AD or PD spectrum, or healthy aged individuals^22^. The participants were classified into 2 × 2 factorial categories based on cognitive disturbance and movement disorder to identify associations between clinical phenotypes and olfaction, as well as their underlying neural mechanisms. This novel strategy enabled us to identify factor-specific patterns related to the neural correlates of hyposmia, which was ascribed to atrophy of the MTL in cognitive disturbance and to that of the olfactory bulb in movement disorder. Notably, patients having both cognitive disturbance and movement disorder showed atrophy of both, resulting in the most severe hyposmia. MTL-GMV and OBV, together with age, can account for approximately 40% of olfaction. Using network modelling, we gained a holistic and causal view of olfactory functioning in AD and PD spectra. Our novel approach enhances our understanding of olfactory impairment in neurodegenerative diseases, highlighting olfactory tests with MTL-GMV and OBV measurement as a non-invasive marker for patient stratification.

Olfaction relies on both peripheral and central machinery. From the periphery, the olfactory bulbs serve as the gateway to the central nervous system for odour processing^42^. Atrophy of the olfactory bulbs has been associated with olfactory dysfunction, as observed in healthy individuals across the lifespan^43^. We developed an original AI model to measure OBV from structural MRIs, which successfully confirmed that OBV indexes olfactory functions. The present study is the first to link olfactory dysfunctions with OBV atrophy in a relatively large-scale patient cohort. In the brain, our results have shown that olfactory function is subserved by the MTL, including the amygdala, olfactory cortex, hippocampus, and parahippocampal gyrus encompassing the entorhinal cortex. The amygdala is a key region involved in higher-order olfactory perception^44^ and olfactory memory^45^, receiving afferents from the olfactory bulb via the piriform and periamygdaloid cortex^46^. The parahippocampal gyrus plays a crucial role in linking memory retrieval with olfactory identification^47^. The hippocampus constitutes the secondary olfactory regions^48^, and smaller hippocampal GMV correlates with poor odour identification^11,49^. Normal aging accompanies progressive olfactory dysfunction, likely reflecting senescent changes within the central olfactory system, including OBV shrinkage^43^ and MTL atrophy^50^. Importantly, these senescence-related structural changes in the olfactory pathways are more pronounced in neurodegenerative processes with disease-specific patterns, as discussed below. Additionally, the OSIT-J score was correlated with MCC-GMV. Structural evidence linking the MCC to olfaction has not been shown before. Although not traditionally considered a core olfactory region, the MCC may modulate olfaction through attentional and affective mechanisms^51^, as supported by MCC activation during odour-emotion tasks^52^. Overall, we identified the olfactory bulbs, MTL, and MCC as the structural correlates of olfactory functions.

Olfactory deficits are associated with an increased risk of cognitive decline^53^ and AD^54^. Our factorial analysis revealed the MTL underlying hyposmia in cognitive disturbance. The cognition-disturbed (C+M− and C+M+) groups showed Aβ deposition, supporting the involvement of AD pathology in their cognitive disturbance. The MTL serves as a hub connecting olfaction with cognition in AD^55^. The MTL—particularly the hippocampus, parahippocampal gyrus, and amygdala—are among the structures most strongly affected by tau pathology in AD.

Tau accumulation is the most reliable molecular indicator of clinical progression in AD^56^, and accumulating evidence shows that cortical atrophy closely mirrors tau pathology^57^. As AD pathology progresses, the distribution of NFTs defines Braak’s three stages: the transentorhinal, limbic, and neocortical ones^58^. NFTs appear very early in the entorhinal and transentorhinal areas, and then in the anterior olfactory nucleus and the olfactory bulb at later stages^19,59–61^. The early tau pathology, involving the entorhinal and transentorhinal areas, likely plays a critical part in mediating coexisting olfactory deficits and cognitive decline in the AD spectrum. Conversely, we failed to find MTL atrophy in the movement disorder-only (C−M+) group, which corresponds to PD with normal cognition. This is reasonable because the MTL is affected only at a late stage in PD^62^, as will be discussed later. Together, we identified the combination of hyposmia and MTL atrophy in the AD spectrum, but not in PD with normal cognition. Olfactory dysfunction coupled with MTL atrophy can be taken as a marker of the AD spectrum.

We confirmed olfactory dysfunction in movement disorders^41^. The movement disorder (C−M+ and C+M+) groups exhibited reduced striatal dopamine transporter binding, consistent with the pathology of PD. Despite the profound olfactory dysfunction, the C−M+ group did not exhibit atrophy in the brain, including the MTL. However, our factorial analysis of AI-computed OBV uncovered abnormalities of the olfactory bulb underlying hyposmia associated with movement disorders. This result agrees with previous small-scale studies that reported reduced OBV in PD compared with healthy controls and individuals with atypical Parkinsonian syndromes^63,64^. PD pathology involving the olfactory pathway follows a reversed pattern from that of AD. According to the Braak staging^62^, Lewy body pathology initially involves the olfactory bulb and the dorsal vagal nucleus. Early involvement of the olfactory bulb may contribute to the early olfactory dysfunctions in PD^65^. In AD, tau pathology can be observed in the olfactory bulbs only at a late stage; moreover, it remains to be demonstrated whether the tau in the olfactory bulb has a significant impact on olfactory impairment^66^. Together, it is plausible that Lewy body pathology in the olfactory bulb uniquely contributes to early olfactory dysfunction in the PD spectrum. Together, the coupling of olfactory dysfunction and OBV reduction suggests Lewy body pathology, which may predict the progression of clinical symptoms from early olfactory impairment to subsequent movement disorders.

We observed the most profound olfactory impairment in the group having both cognitive disturbance and movement disorder (C+M+). Previous studies have reported an association between poor olfaction and cognitive disturbance in PD^67^ or with cognitive complaints and slow gait in a community cohort^68^; however, the neural correlates of such associations have not been elucidated to date. The C+M+ group had pronounced atrophy of MTL, accompanied by moderate shrinkage of the olfactory bulbs. The severe olfactory dysfunction mainly reflected the additive influence of abnormalities involving both. This “double hit” pattern of structural damage very likely resulted in the most severe olfactory dysfunction among the four groups. This theory is underpinned by stepwise linear regression analysis, demonstrating that OBV and MTL-GMV both contribute to olfaction as independent variables. These observations also suggest a complex and multifactorial pathomechanism underlying hyposmia in the C+M+ group. Our preliminary analysis suggests that AD-PD co-pathology existed in the subgroup of C+M+ with relatively high Aβ deposition, whereas the remaining C+M+ subgroup had more advanced PD pathology (data not shown). This finding will be formally addressed using the complete dataset of the PADNI cohort. Overall, we propose that the C+M+ group presenting with the most severe hyposmia had the “double hit” pathology involving both the MTL and the olfactory bulbs. This discovery has significantly advanced our clinically feasible stratification strategy of neurodegenerative diseases spanning the AD and PD spectra.

Our SEM analysis provided a comprehensive view of the relationship across the demographic, behavioural, and neural correlates underlying hyposmia. The SEM identified two processes contributing to olfactory impairment: an efferent process from the MTL and an afferent process from the olfactory bulbs. In the AD spectrum, MTL atrophy impairs olfactory discrimination required for OSIT-J partly in relation to cognitive decline, consistent with prior evidence linking hippocampal-entorhinal degeneration to odour identification^69^. In the PD spectrum, OBV reduction was directly related to olfactory dysfunction, which was correlated with motor impairment. While this correlation does not necessarily imply a direct causal link, it suggests that olfactory bulb atrophy and motor impairments may progress in parallel as manifestations of common α-synuclein pathology^70^. Notably, age exerted an additional influence on both MTL and OBV atrophy, potentially amplifying olfactory dysfunctions due to pathological changes^43,71^. These results suggest that considering the disease-specific, directional process of propagation is essential to utilize hyposmia as a clinically relevant marker across neurodegenerative disorders.

This study has several limitations. First, the sample size within each symptom-based subgroup (e.g., C+M+) was modest, which may reduce the statistical power to detect subtle group differences or interaction effects. For example, we excluded prodromal AD/PD due to the limited number of participants. We will address this issue with larger, more balanced samples to address the related issues in prodromal conditions in future reports. The observed residual correlations between olfactory performance and both MoCA-J and UPDRS-III scores suggest that non-structural factors may contribute to olfactory decline, highlighting the need for multimodal approaches, including functional connectivity data, in future multimodal data analysis. Finally, although our cohort includes longitudinal follow-up, the present analysis was restricted to baseline cross-sectional data. Future studies using the longitudinal dataset with longer follow-up will be essential to directly address the directionality of propagation.

We propose that two major epicentres, the MTL and olfactory bulbs within the olfactory pathway, underlie hyposmia in the two major neurodegenerative entities, the AD and PD spectra. The double hit of the MTL and the olfactory bulbs may result in the most severe olfactory dysfunction, as observed in patients with both cognitive disturbance and movement disorder. We further propose that it is essential to consider the directionality of pathomechanism propagation: the efferent process from MTL and the afferent process from the olfactory bulbs. The scrutiny of olfactory impairment provides a unique path toward an understanding of pathomechanisms of the AD and PD spectra, offering guidance for early diagnosis and refined therapeutic strategies.

## Data availability

Data used in this study were obtained from the PADNI database. These data are not publicly available due to restrictions on participant privacy and consent. Still, access may be granted to qualified researchers upon reasonable request and subject to approval by the PADNI data access committee.

