## supplementary materials for "Neural Underpinnings of Olfactory Dysfunction across Parkinson’s and Alzheimer’s Spectra"

### Supplemental Methods

#### S1 ComBat Harmonization for VBM analysis

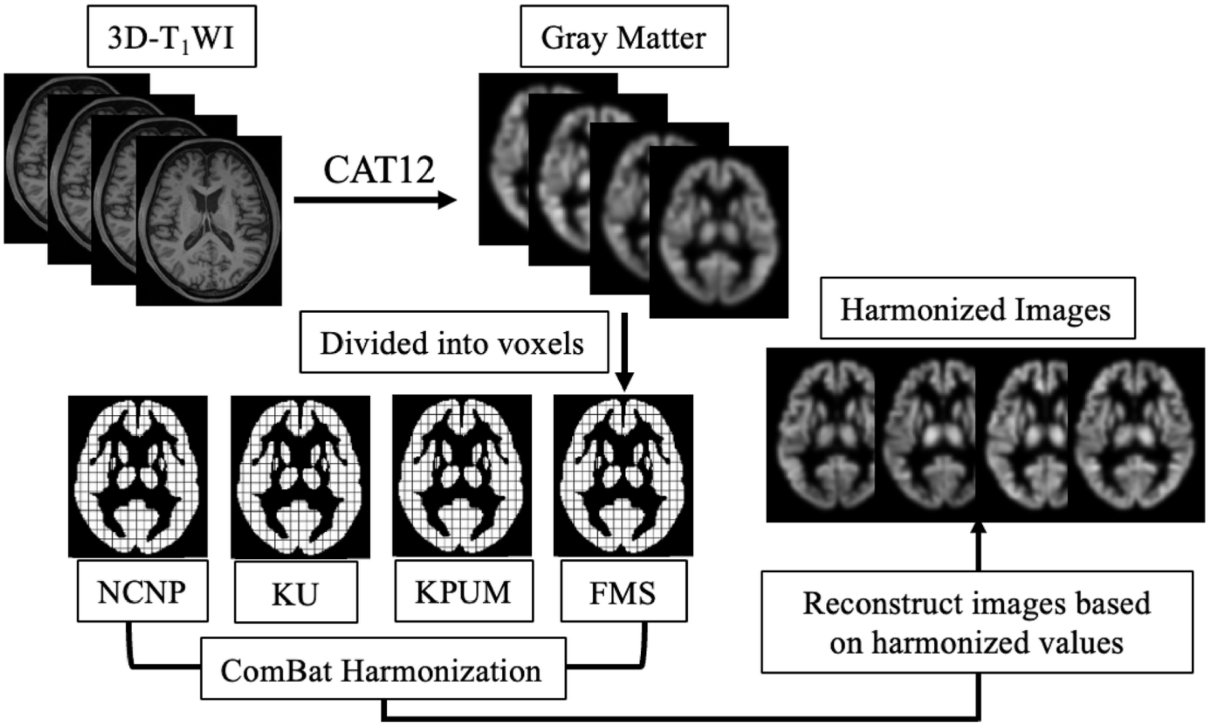


**Figure 1. Flowchart of ComBat Harmonization.**

In the present study, T1-weighted MRIs were collected from four different centers. We performed ComBat harmonization to mitigate the site effects. ComBat operates as an empirical Bayes-based method designed to minimize differences (mean and variance) between data, including inter-site and scanner variations[1]. Specifically, we used the neuroCombat implementation developed by Fortin (available at: https://github.com/Jfortin1/ComBatHarmonization), which has been successfully employed in prior neuroimaging studies to harmonize diffusion tensor imaging and cortical thickness data[2,3]. ComBat has been extended to voxel-wise applications in gray matter morphometry studies[4]. We applied the voxel-wise ComBat method to the original value at each voxel in the gray matter images after the spatial normalization process and subsequently generated new NIfTI images based on the ComBat-harmonized values (Figure 1). The imaging sites and the scanners were chosen as the variables to be removed from the data, whereas the biological variables (age, gender, TIV, group label, MoCA-J, UPDRS-III, and OSIT-J) were modeled to be retained.

To evaluate the effectiveness of the ComBat adjustment, we performed repeated-measures ANCOVA (RM-ANCOVA) with age, gender, TIV, OSIT-J score, and disease labels as covariates, including a within-subject factor harm (levels: before vs. after ComBat). The analysis revealed a systematic shift in MTL values (harm: $F(1,212) = 26.93$, $p < 0.001$), having significant interactions with the site factor ($F(3,212) = 9.15$, $p < 0.001$). Separate ANCOVAs on the MTL-GMV values before and after ComBat confirmed that the main effect of site was reduced after ComBat (pre: $F(1,214) = 0.14$, $p = 0.71$, partial $\eta^2 = 0.00065$; post: $F(1,214) = 0.03$, $p = 0.86$, partial $\eta^2 = 0.000056$). Figure S2 exemplifies MTL-GMV (comparing C−M− and C+M− groups only for brevity) to further illustrate the site-related effects and impact of ComBat harmonization. In the multi-site data, the effect size of the multi-site data became larger after ComBat ($d = -1.71$) compared to before ComBat ($d = -1.64$).

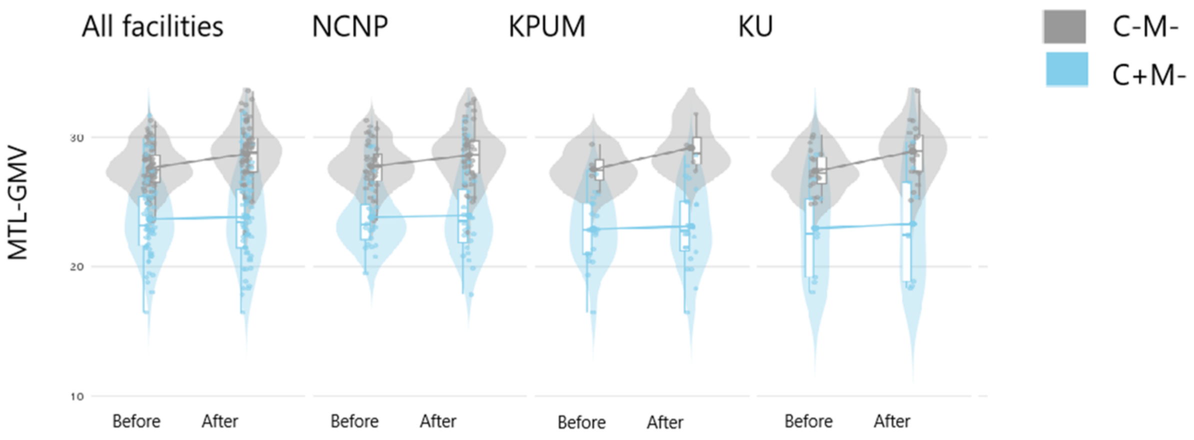


**Figure 2. MTL (cognition-related: C−M−, C+M−) distributions across sites before and after ComBat.** Results showing reduced inter-site variability with preserved group differences. Note that the group comparison at FMU is not shown because the C−M− data were unavailable. NCNP, National Center of Neurology and Psychiatry; KPUM, Kyoto Prefectural University of Medicine; KU, Kyoto University; FMU, Fukushima Medical University.

### S2 Olfactory Bulb Segmentation Model

#### Image processing and annotation

Sixty participants were randomly selected from the interim PADNI database, and their data were used as the derivation dataset for developing the prediction model. In addition to the derivation dataset, ten participants were selected from each of the four groups for internal validation of the segmentation model's accuracy. For these images, we performed manual segmentation for the annotation of the olfactory bulb, using ITK-SNAP (version 4.0.2). The olfactory bulb is an almond- or spindle-shaped, bilaterally symmetric structure located ventral to the orbitofrontal cortex[5]. On T2-weighted MRIs, the olfactory bulb appears as a T2 low-intensity structure surrounded by a high-intensity signal of cerebrospinal fluid, providing a clear contour[6]. T2-weighted images in these datasets were cropped to 32 voxels from the centroid of the annotated olfactory bulbs (100 pairs).

For the MRIs of other participants, the location of the olfactory bulb was first manually annotated on the MNI standard template. This olfactory bulb reference was transformed to the native space of each participant using the inverse deformation field from the normalization step of preprocessing. The referential location for cropping was subsequently calculated, and the T2-weighted images were cropped using the center of the reference as the center. Thus, T2-weighted images of all participants were cropped to a size of 64 × 64 × 64 voxels at the center of the olfactory bulb. The voxel values were standardized using the mean and standard deviation calculated from voxel intensities within the field of view, after removing the 1% of values with the lowest and highest intensities.

#### Constructing the olfactory bulb prediction model

We found it suboptimal just to apply a previously published method for automated olfactory bulb segmentation on T2-weighted MRI[7] to our dataset (Figure S4), yielding a mean Dice coefficient of 0.601±0.173. This finding prompted us to develop a customized segmentation model tailored to our T2-weighted MRI data.

A predictive model based on deep learning was constructed and trained using the annotated data from the 60 participants in the derivation cohort as supervisory information. The model was constructed using Python 3.8 with PyTorch (version 2.4.1, CUDA 12.1 build), with a 3D U-Net as the basic structure[8]. Instead of conventional convolutional blocks, residual blocks were employed throughout both the encoder and decoder pathways to improve gradient flow and mitigate vanishing gradients. The activation function used in all residual blocks was the leaky rectified linear unit, with a negative slope coefficient of 0.2, following the configuration described in a previous study[9]. The model was trained for 100 epochs using a combined loss function consisting of Dice loss and binary cross-entropy loss (weighted 0.7 and 0.3, respectively). Optimization was performed using the AdamW optimizer with an initial learning rate of $1.0 \times 10^{-4}$ and a weight decay of $1.0 \times 10^{-5}$, following the decoupled weight decay strategy proposed by Loshchilov and Hutter[10]. A ReduceLROnPlateau scheduler was applied to dynamically adjust the learning rate. The rate was halved if the validation loss did not improve over 10 consecutive epochs, with a minimum learning rate threshold of $1.0 \times 10^{-6}$. The data was augmented through three-dimensional image transformation using scaling, translation, rotation, flipping, and the addition of Gaussian noise. Each image of the training dataset was randomly transformed by applying these three transformations using a linear interpolation algorithm, including random flipping along the width axis ($p = 0.5$), random rotation within ± 5°, translation within ±3 voxels, scaling within 90 – 110%, and addition of Gaussian noise (std: 0.01 – 0.05, $p = 0.5$). All computations were performed on a workstation equipped with an NVIDIA RTX 6000 Ada Generation GPU (48 GB VRAM). Model generalizability was assessed via five-fold cross-validation within the derivation cohort. For the final inference, segmentation predictions were generated by ensembling the best models trained across all 5 folds. The network's final output passed through a sigmoid activation to obtain voxel-wise probability maps, which were then binarized using a threshold of 0.5 to generate the final segmentation masks. We inspected the anatomical consistency of the results; all segmented regions were consistent with anatomical structures, and no apparent isolated noise-like fragments were observed. Consequently, no additional morphological post-processing was performed.

#### ComBat Harmonization for OBV

Similarly, to mitigate potential site effects, we also applied ComBat harmonization to the multisite OBV data. ANCOVAs performed separately for pre- and post-ComBat values showed that the site effect was insignificant even before ComBat (pre: $F(1,214) = 0.006$, $p = 0.94$; post: $F(1,214) = 0.007$, $p = 0.93$). Figure S5 displays the total OBV between the C−M− and C−M+ groups to illustrate that the site effects were negligible for OBV. For all facilities, the total OBV was significantly lower in C−M+ group compared to C−M− group both before and after ComBat, with stable effect size before (d = −0.78) and after ComBat (d = −0.78), indicating that harmonization reduced inter-site variability without attenuating the group differences, thereby supporting the robustness of the olfactory bulb automatic segmentation model across facilities.

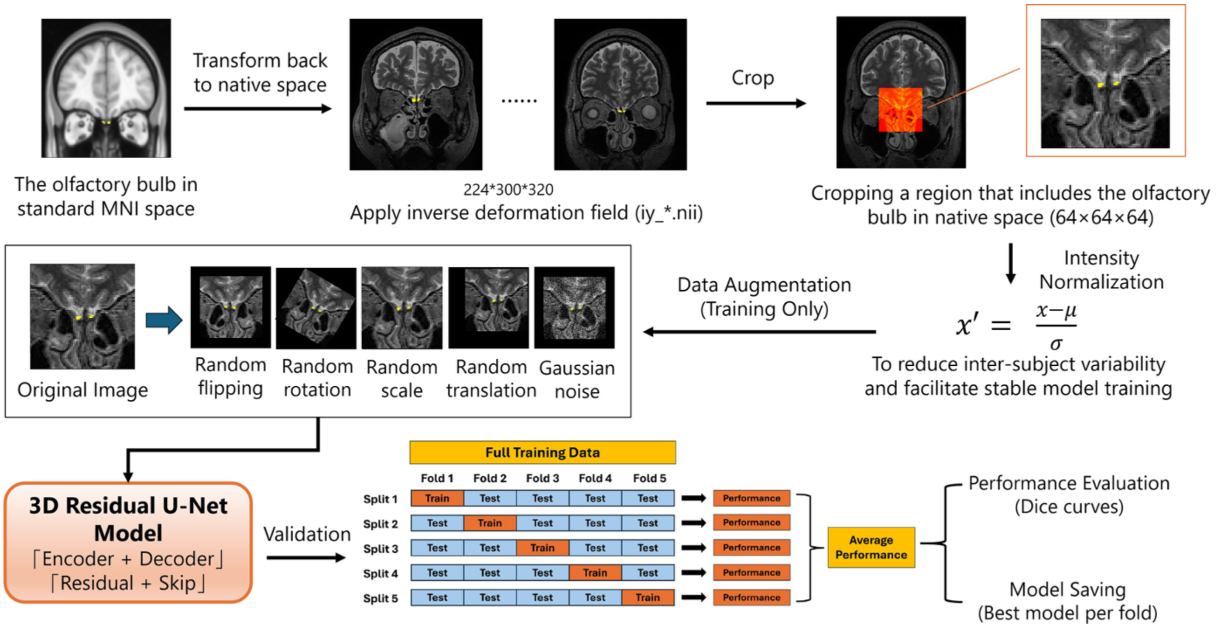


**Figure 3. Deep Learning-Based Workflow for Olfactory Bulb Segmentation**

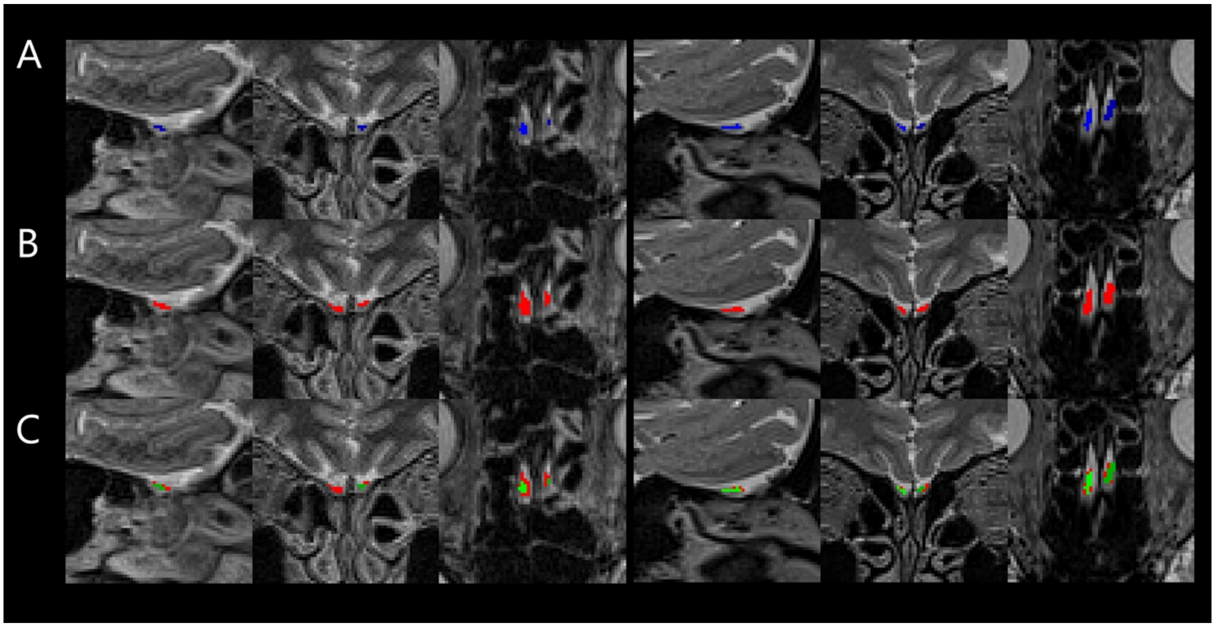


**Figure 4. Comparison of olfactory bulb segmentation methods.** (**A**) Olfactory bulb from the previously published olfactory bulb segmentation method (blue); (**B**) Olfactory bulb from the

customized segmentation model trained on our database (red); (**C**) An overlay comparison indicating that the previous segmentation model (green) showed lower accuracy than our customized segmentation model (red).

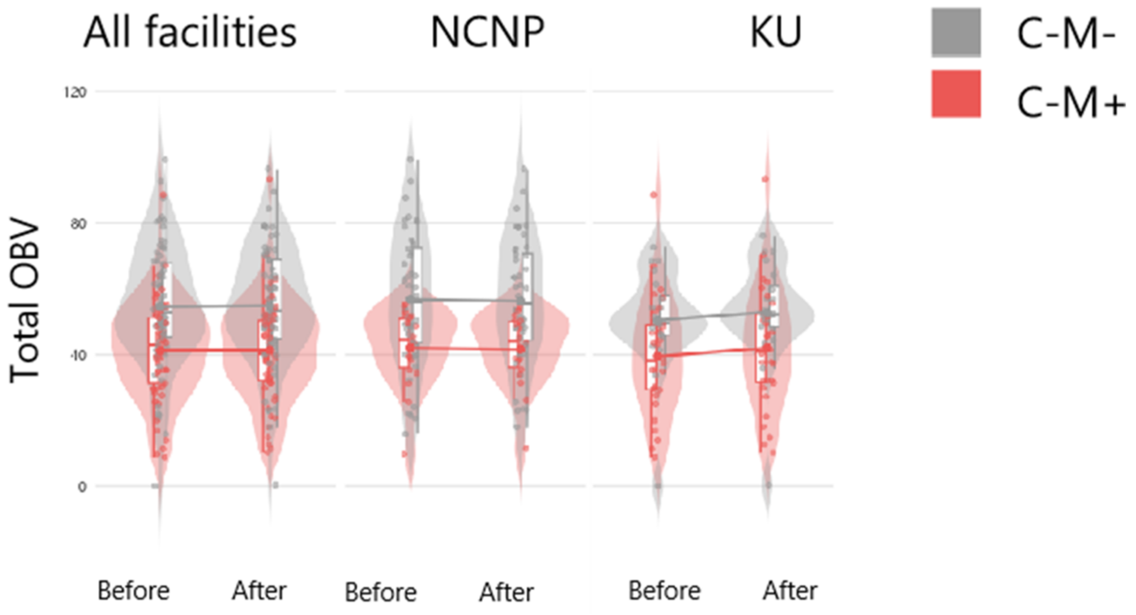


**Figure 5. Total OBV (motor-related: C−M−, C−M+) distributions across sites before and after ComBat.** Results showing reduced inter-site variability with preserved group differences. Note that C−M+ data were unavailable at KPUM, and C−M− data were unavailable at FMU (graphs not shown). NCNP, National Center of Neurology and Psychiatry; KPUM, Kyoto Prefectural University of Medicine; KU, Kyoto University; FMU, Fukushima Medical University.

### Supplementary Results

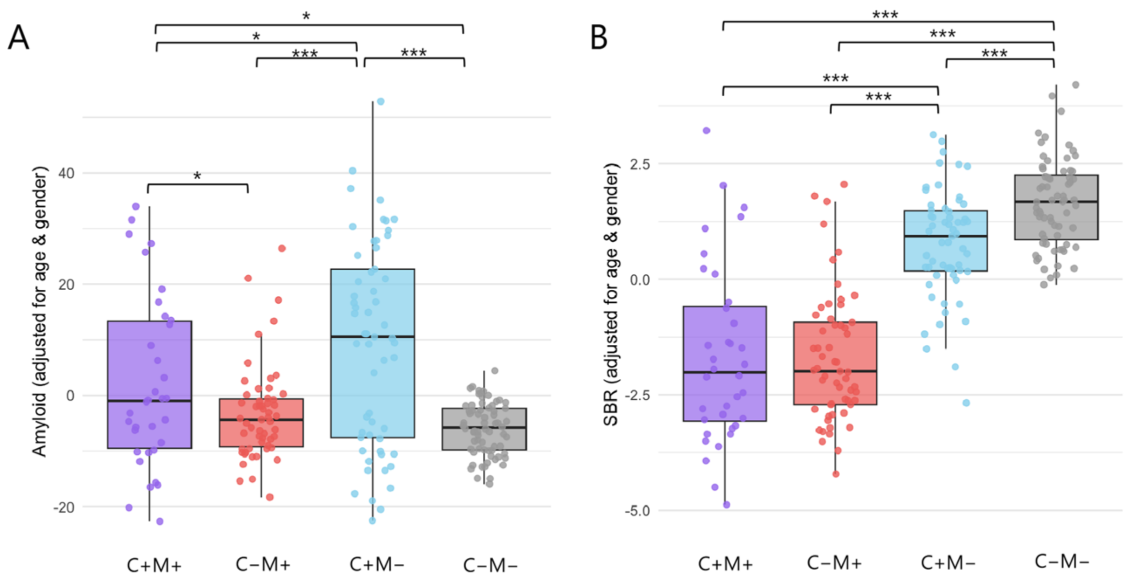


**Figure 6: Comparison of molecular markers across groups.** Age and gender were included as covariates in the analysis. (**A**) Comparison of Amyloid-β load (AβL) derived from amyloid-PET across groups[11]. The groups with cognitive impairment (C+M+ and C+M−) exhibited higher amyloid burden compared to cognitively normal groups (C−M+ and C−M−), with the cognitive impairment-only group (C+M−) showing the highest level of amyloid burden among all groups. (**B**) Comparison of bilateral mean Specific Binding Ratio (SBR) derived from dopamine transporter single-photon emission computed tomography (DAT-SPECT). The groups with motor impairment (C+M− and C−M+) exhibited lower SBR compared to motor normal groups (C+M− and C−M−), with the cognitive impairment-only group (C+M−) exhibiting lower SBR compared to the symptom-free individuals. Significance levels are denoted as follows: $p < 0.05$ (*), $p < 0.001$ (***). Note: Amyloid-PET data were unavailable for 8 participants, and DAT-SPECT data were unavailable for 2 participants.

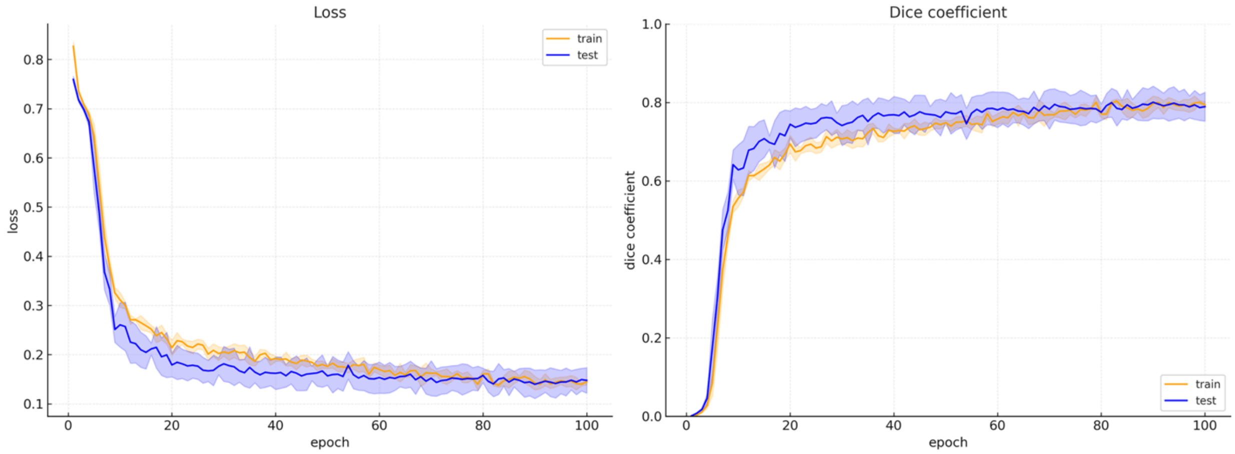


**Figure 7: Training and validation performance of the residual 3D U-Net model.** The left panel shows the training and validation loss curves over 100 epochs, demonstrating stable convergence with decreasing loss. The right panel displays the Dice coefficient curves, indicating a consistent improvement in segmentation accuracy. The final validation Dice coefficient reached 0.8059 ± 0.0378, suggesting good generalization performance of the model.

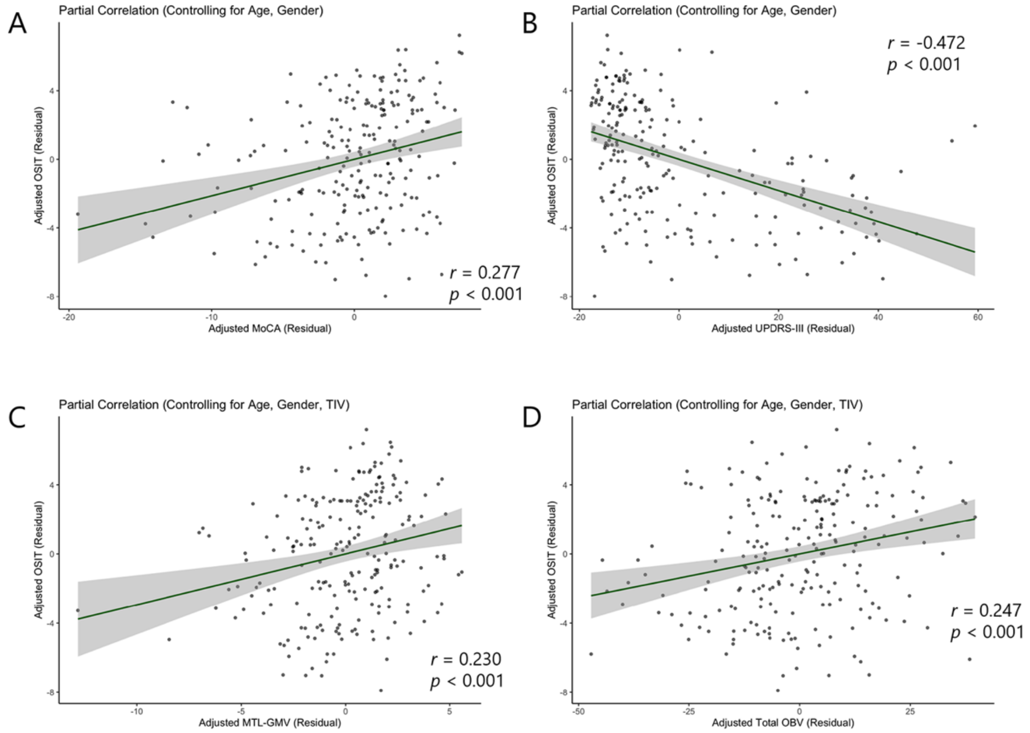


**Figure 8. Partial correlations of olfactory function (OSIT-J) with cognitive performance, motor symptoms, brain volume, and olfactory bulb volume.** (**A**) Partial correlation between MoCA-J and OSIT-J, r=0.277, p<0.001, age and gender were included as covariates in the analysis;

(**B**) Partial correlation between UPDRS-III and OSIT-J, r=-0.472, p<0.00, age and gender were included as covariates in the analysis; (**C**) Partial correlation between OSIT-J and MTL-GMV, r=0.230, p<0.001, age, gender and gender were included as covariates in the analysis; (**D**) Partial correlation between OSIT-J and total OBV, r=0.247, p<0.001, age, gender and gender were included as covariates in the analysis. Abbreviations: UPDRS-III, Unified Parkinson's Disease Rating Scale Part III; MoCA-J, the Japanese version of the Montreal Cognitive Assessment; OSIT-J, Odor Stick Identification Test for Japanese; MTL-GMV, gray matter volume of the middle temporal lobe; OBV, olfactory bulb volume.

**Table 1 Factorial ANCOVA of MoCA-J scores with age as covariate**

| Effect | df | F | p-value | Significance |
|---|---|---|---|---|
| C factor | 1, 217 | 121.66 | < 0.001 | *** |
| M factor | 1, 217 | 2.22 | 0.137 | ns |
| Age (cov) | 1, 217 | 6.95 | 0.009 | ** |
| C × M | 1, 217 | 11.32 | < 0.001 | *** |

Note: Results from factorial ANCOVA on MoCA-J scores with age included as a covariate. Effects of cognitive status, motor status, and their interaction are reported. df = degrees of freedom; ns = not significant. Significance levels: $p < 0.01$ (**), $p < 0.001$ (***).

**Table 2 Post hoc comparisons of OSIT-J scores across the four groups**

| Contrast | Estimate | Std. Error | t value | *p*-value | Significance |
|---|---|---|---|---|---|
| C−M+ vs. C+M+ | 0.7058 | 0.6203 | 1.138 | 0.66485 | n.s. |
| C+M− vs. C+M+ | 2.2105 | 0.5921 | 3.734 | 0.00131 | ** |
| C−M− vs. C+M+ | 5.0319 | 0.6064 | 8.298 | <0.001 | *** |
| C+M− vs. C−M+ | 1.5046 | 0.5512 | 2.730 | 0.03451 | * |
| C−M− vs. C−M+ | 4.3261 | 0.5056 | 8.556 | <0.001 | *** |
| C−M− vs. C+M− | 2.8215 | 0.5364 | 5.260 | <0.001 | *** |

Note: Significance levels are denoted as follows: $p < 0.05$ (*), $p < 0.01$ (**), $p < 0.001$ (***). Abbreviations: C+M+, cognitive impairment with movement disorder; C–M+, cognitive normal with movement disorder; C+M–, cognitive impairment with motor normal; C–M–, cognitive normal with motor normal.

**Table 3 Post hoc comparisons of MTL-GMV across the four groups**

| Contrast | Estimate | Std. Error | t value | Adjusted p-value | Significance |
|---|---|---|---|---|---|
| C−M+ vs. C+M+ | 0.27379 | 0.07306 | 3.747 | 0.00127 | ** |
| C+M− vs. C+M+ | -0.14778 | 0.06964 | -2.122 | 0.14855 | n.s. |
| C−M− vs. C+M+ | 0.42027 | 0.07133 | 5.891 | <0.001 | *** |
| C+M− vs. C−M+ | -0.42157 | 0.06481 | -6.505 | < 0.001 | *** |
| C−M− vs. C−M+ | 0.14648 | 0.05998 | 2.442 | 0.07174 | n.s. |
| C−M− vs. C+M− | 0.56805 | 0.06345 | 8.952 | <0.001 | *** |

Note: Significance levels are denoted as follows: $p < 0.01$ (**), $p < 0.001$ (***). Abbreviations: C+M+, cognitive impairment with movement disorder; C–M+, cognitive normal with movement disorder; C+M–, cognitive impairment with motor normal; C–M–, cognitive normal with motor normal.

**Table 4 *Post hoc* comparisons of mean MCC-GMV across the four groups**

| Contrast | Estimate | Std. Error | t value | Adjusted p-value | Significance |
|---|---|---|---|---|---|
| C−M+ vs. C+M+ | 0.04689 | 0.01331 | 3.522 | 0.0029 | ** |
| C+M− vs. C+M+ | -0.01504 | 0.01269 | -1.185 | 0.6351 | n.s. |
| C−M− vs. C+M+ | 0.07021 | 0.01300 | 5.402 | <0.001 | *** |
| C+M− vs. C−M+ | -0.06193 | 0.01181 | -5.245 | < 0.001 | *** |

| | | | | | |
|---|---|---|---|---|---|
| C−M− vs. C−M+ | 0.02332 | 0.01093 | 2.134 | 0.1446 | n.s. |
| C−M− vs. C+M− | 0.08525 | 0.01156 | 7.374 | <0.001 | *** |

Note: Significance levels are denoted as follows: $p < 0.01$ (**), $p < 0.001$ (***). Abbreviations: C+M+, cognitive impairment with movement disorder; C–M+, cognitive normal with movement disorder; C+M–, cognitive impairment with motor normal; C–M–, cognitive normal with motor normal.

**Table 5 Comparison of model fit indices for structural equation models.**

| Model | Parameters | $\chi^2$ (df), p | CFI | TLI | RMSEA (90% CI) | SRMR | AIC | BIC | Note |
|---|---|---|---|---|---|---|---|---|---|
| Final model | 19 | 7.938 (6), *p*=0.243 | 0.994 | 0.981 | 0.038 (0–0.101) | 0.032 | 2828 | 2892 | Dual-pathway model |
| Alternative model | 20 | 7.930 (5), *p*=0.160 | 0.991 | 0.966 | 0.051 (0–0.115) | 0.031 | 2830 | 2898 | Dual-pathway model +OBV–MTL covariance |

Note: Model fit indices for the final SEM model and an alternative model including an OBV–MTL covariance. The added path was not significant ($r = 0.005$, $p = 0.093$) and worsened fit, supporting the independence of top-down and bottom-up pathways.
